# Cdc42 activation is necessary for heterosynaptic cooperation and competition

**DOI:** 10.1101/2023.09.26.559525

**Authors:** Mariana Nunes, Natália Madeira, Rosalina Fonseca

## Abstract

Synapses change their weights in response to neuronal activity and in turn, neuronal networks alter their response properties and ultimately allow the brain to store information as memories. As for memories, not all events are maintained over time. Maintenance of synaptic plasticity depends on the interplay between functional changes at synapses and the synthesis of plasticity-related proteins that are involved in stabilizing the initial functional changes. Different forms of synaptic plasticity coexist in time and across the neuronal dendritic area. Thus, homosynaptic plasticity refers to activity-dependent synaptic modifications that are input-specific, whereas heterosynaptic plasticity relates to changes in non-activated synapses. Heterosynaptic forms of plasticity, such as synaptic cooperation and competition allow neurons to integrate events that occur separated by relatively large time windows, up to one hour. Here, we show that activation of Cdc42, a Rho GTPase that regulates actin cytoskeleton dynamics, is necessary for the maintenance of long-term potentiation (LTP) in a time-dependent manner. Inhibiting Cdc42 activation does not alter the time-course of LTP induction and its initial expression but blocks its late maintenance. We show that Cdc42 activation is involved in the phosphorylation of cofilin, a protein involved in modulating actin filaments and that weak and strong synaptic activation leads to similar levels on cofilin phosphorylation, despite different levels of LTP expression. We show that Cdc42 activation is required for synapses to interact by cooperation or competition, supporting the hypothesis that modulation of the actin cytoskeleton provides an activity-dependent and time-restricted permissive state of synapses allowing synaptic plasticity to occur. We found that under competition, the sequence in which synapses are activated determines the degree of LTP destabilization, demonstrating that competition is an active destabilization process. Taken together, we show that a dynamic actin cytoskeleton is necessary for the expression of homosynaptic and heterosynaptic forms of plasticity. Determining the temporal and spatial rules that determine whether synapses cooperate or compete will allow us to understand how memories are associated.

**Graphical Abstract:** 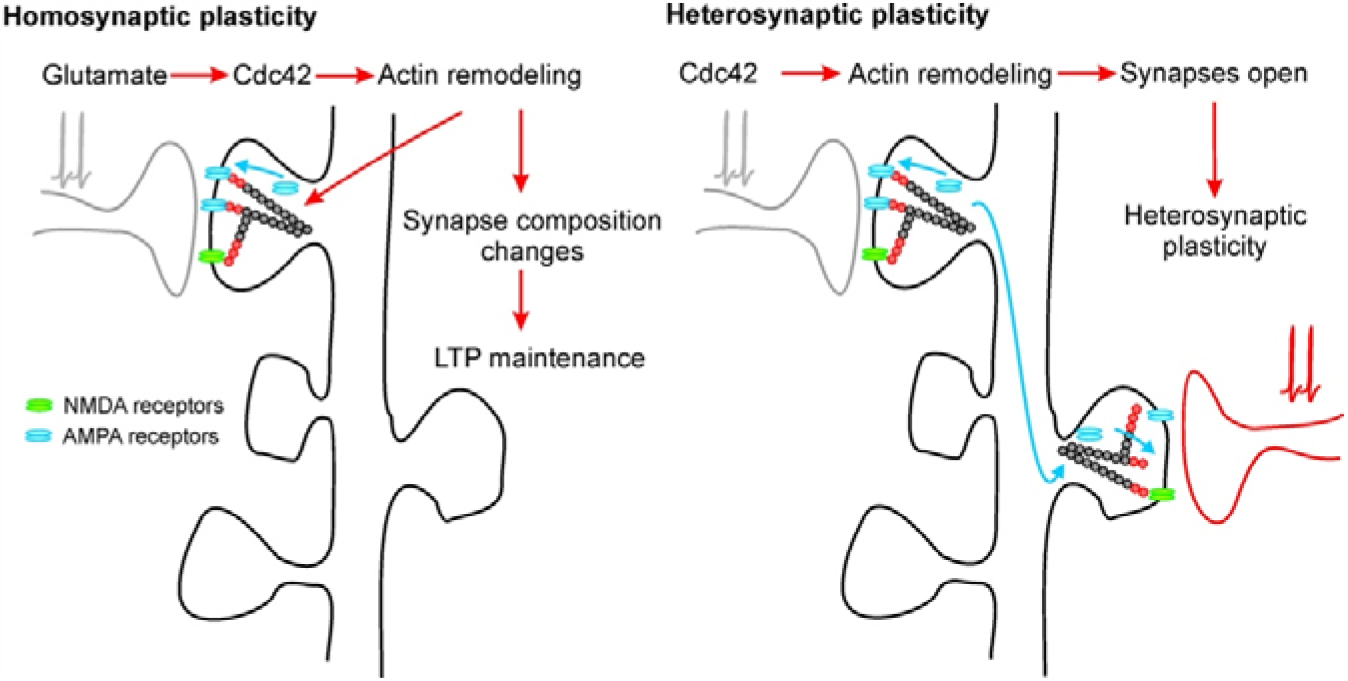

**Highlights:**

- Cdc42 is required for the maintenance of homosynaptic synaptic plasticity
- Weak and strong stimulation modulate actin by cofilin phosphorylation
- Cdc42 activation is necessary for heterosynaptic cooperation and competition
- Synaptic competition is an active destabilization process
- The time-window of synaptic cooperation and competition is activity dependent

## Introduction

It is widely accepted that modifications in the strength of connections between neurons are a cellular mechanism through which the brain encodes and storages information (Mayford et al., 2012). Changes in synapses reflect a combination of functional and structural alterations at synapses, including long-term potentiation (LTP) and depression (LTD) as well as morphological plasticity of dendritic spines (Martin et al., 2000). LTP has many interesting properties that make it an attractive cellular model of circuitry refinement and learning and memory, such as cooperativity, associativity, input-specificity and persistence. The persistence of LTP is thought to be dependent on the synthesis of plasticity-related proteins (PRPs), which stabilizes the functional changes induced by neuronal activity (Fonseca et al., 2006; Mayford et al., 2012). But how could the stabilization occur in the correct place? Given that synaptic plasticity is, by large, input-specific, then PRPs would be either synthesized locally or available cell-wide and targeted or captured at activated synapses. The synaptic tagging and capture hypothesis (STC) proposes that activated synapses set up molecular tags, which allow synapses to capture newly synthesized PRPs in an input-specific manner (Frey and Morris, 1997; Redondo and Morris, 2011). The STC hypothesis states that the maintenance of synaptic plasticity is based on the interplay of two linked, but independent, cellular processes. Upon the induction of LTP or LTD, an input-specific and time-limited synaptic tag is generated. If the initial stimulus is not strong enough to trigger protein synthesis (weak stimulus), then a transient form of plasticity is expressed. If the stimulus is strong enough (strong stimulus), then protein synthesis is triggered and the subsequent capture of PRPs in activated synapses allows the maintenance of plasticity. Synaptic tagging and the expression of the transient forms of LTP (or LTD) occur without protein synthesis or are insensitive to protein synthesis inhibitors but the maintenance of synaptic changes depends on PRPs capture.

An interesting twist of the experimental demonstration of the STC model is the observation that PRPs can be shared between different sets of synapses. The induction of a transient LTP (weak) set up tags that can capture PRPs synthesized by the strong stimulation of a second independent synaptic input (strong). This observation shows that synapses can interact with each other. Homosynaptic plasticity gives rise to a form of heterosynaptic cooperation in which different groups of activated synapses, weak and strong, interact by sharing PRPs and maintaining LTP. Interestingly, synapses engage in competition if the number of activated inputs increases creating a competitive pressure that destabilizes LTP maintenance (Fonseca et al., 2004; Sajikumar et al., 2014). In the light of the STC, the idea behind synaptic competition is that all activated synapses (weak and strong) will be able to capture PRPs from the common pool creating a competitive pressure that will interfere with LTP maintenance. Synaptic cooperation and competition are therefore two forms of heterosynaptic plasticity based on the interaction of biochemical signals that allow synapses to integrate cell wide activity within large time windows (Pinho et al., 2020; Szabó et al., 2016).

If one considers the nature of the synaptic tag, one strong candidate is the actin cytoskeleton. It is well established that morphological changes observed in dendritic spines, induced by synaptic plasticity, are strongly associated with the actin cytoskeleton dynamics (Honkura et al., 2008; Matsuzaki et al., 2004). The dynamics of actin is modulated by several actin-binding proteins (ABPs) such as CaMKII, Cofilin, and Drebrin (Hayashi et al., 1996; Okamoto et al., 2009, 2004). These actin regulators are activated/inactivated in an activity-dependent manner, modulating the rate of F-actin assembly, disassembly, stabilization, and bundling during synaptic plasticity (Pinho et al., 2020). Previous results from our laboratory have indeed shown a critical role of actin dynamics in the tagging and capture of PRPs in both LTP and LTD, supporting the hypothesis that an activity-dependent remodeling of F-actin, through CaMKII activation, renders the synapse locally and transiently permissive to plasticity modifications (Fonseca, 2012; Szabó et al., 2016). Interestingly, CaMKII leads to activation of Cdc42, a Rho GTPase that also plays a role in regulating the actin cytoskeleton in dendritic spines (Kim et al., 2014, 2015). Cdc42 is known to be upstream of PAK (p21-activated kinase), which phosphorylates and activates LIM-kinase (LIMK), which, in turn, phosphorylates cofilin, inhibiting its actin-depolymerizing activity (Hodge and Ridley, 2016; Kim et al., 2014; Sumi et al., 1999). Cdc42 is also responsible for interacting with the WAVE1/N-WASP pathway to activate Arp2/3 complex-dependent actin polymerization. Thus, Cdc42 activation promotes actin polymerization by indirectly increasing cofilin phosphorylation. Since Cdc42 is activated by synaptic plasticity in a time-resricted manner and its activity is heavily restricted to the stimulated spine (input-specificity) (Murakoshi et al., 2011), we hypothesize that Cdc42 plays a crucial role in the setting of the synaptic tag, promoting input-specific maintenance of synaptic plasticity. In this work, we tested whether inhibiting the activation of Cdc42 interferes with the maintenance of both homosynaptic and heterosynaptic forms of LTP.

## Methods

### Slice preparation

All experiments were performed in transverse hippocampal slices from weaned male Wistar Han rats (P21-P35), bred at the housing facility of the host institution (NOVA Medical School – Lisbon, Portugal). The procedures were approved by the Portuguese National Authority for Animal Health (DGAV) and the Ethical Committee of the NOVA Medical School and are in accordance with the Decree-Law No. 113/2013 of 7 August (based on the EU Directive No. 2010/63 on the protection of animals used for scientific or educational purposes). Animals were decapitated under isoflurane anesthesia and the brains were quickly removed and immersed in ice-cold artificial cutting cerebro-spinal fluid (ACSF). Cutting ACSF was saturated with 95%O_2_/5%CO_2_ and contained (in mM): NaCl 126, KCl 2.5, NaH_2_PO_4_ 1.25, NaHCO_3_ 26, MgCl_2_ 5, CaCl_2_ 1, Glucose 25. The hippocampi were isolated and cut into 400 μm-thick transverse slices using either a vibrotome (Leica, VT1200S) or a tissue slicer (Siskiyou, Inc.). Slices were maintained in ACSF at 32º C for at least one hour before recording. They were then transferred to a submersion chamber and perfused continuously (1.5-2 ml/min) with recording ACSF at 32º C. The recording ACSF was saturated with 95%O_2_/5%CO_2_ and contained (in mM): NaCl 126, KCl 2.5, NaH_2_PO_4_ 1.25, NaHCO_3_ 26, MgCl_2_ 2, CaCl_2_ 2.8, Glucose 25.

### Electrophysiological recordings

Recordings started after 20 minutes resting phase in the recording chamber. Schaffer collaterals were stimulated with 0.2 ms pulses using monopolar epoxy-insulated tungsten electrodes (Science Products, GmBH, Germany). Field excitatory postsynaptic potentials (fEPSP) were recorded extracellularly in the *stratum radiatum* of the CA1 region (∼130 μm below the slice surface) using glass microelectrodes filled with 3M NaCl (immobilized with agarose - tip resistance 3-5MΩ). Stimulus intensities were set to evoke 50% of the maximal fEPSP slope. The test pulse frequency is 0.33 Hz. LTP was induced after recording a stable 20 minutes baseline of fEPSPs. For experiments that required the activation of three pathways, a third stimulating electrode was used to stimulate an independent input in the antidromic direction (S3). Input presynaptic independence was assessed by paired-pulse facilitation (PPF) evoked with a 30 ms inter-pulse interval.

### Induction of Long-Term Potentiation

After baseline stimulation, one of the pathways was arbitrarily chosen to receive a tetanus of 25 pulses at a frequency of 100 Hz, repeated 5 times with a 3s interval. The second pathway served as a control pathway and was continuously recorded. In cooperation experiments, a transient form of LTP was induced by stimulation of one pathway with a weak tetanization protocol (Weak-LTP; 25 pulses, 100Hz, repeated 2 times with a 3s interval). After 30 minutes, a maintained form of LTP was induced in a second independent pathway with a strong tetanization protocol (Strong-LTP; 25 pulses, 100Hz, repeated 5 times with a 3s interval). In cooperation experiments, the third stimulated pathway was continuously recorded as a control pathway. Competition was induced by inducing a Weak-LTP in the third input pathway, using the same stimulation parameters as for cooperation experiments (Weak-LTP; 25 pulses, 100Hz, repeated 2 times with a 3s interval; Strong-LTP; 25 pulses, 100Hz, repeated 5 times with a 3s interval).

### Drug treatment

Drugs were dissolved in DMSO and diluted down to achieve the final concentration: ML141 (TargetMol) 5 μM (0.01% DMSO) Cytochalasin-B (TargetMol) 0,5 μM (0.01% DMSO), Jasplakinolide (Invitrogen) 0,1 μM (0.01% DMSO). For the control experiments, only DMSO (0.01%) was added to the ACSF.

### Western-blot analysis of cofilin phosphorylation

Slices of the hippocampus (400μm) were prepared as described for electrophysiological recordings. Slices were placed in the recording chamber and synaptic responses were evoked as described before. After recording baseline responses, LTP was induced by a weak or a strong stimulation and fEPSPs were recorded further for 30 minutes. Slices were then collected and the tissue containing the CA1 region was isolated. The tissue was then deeply frozen in liquid nitrogen and stored at -80°C until further processed. For the western blot analysis of cofilin (Total fraction or phosphorylated fraction), the samples were homogenized with a lysis buffer (0.32M Sucrose, 50 mM Tris Base and protease/phosphatase inhibitor cocktail (Roche, Germany) using a sonicator (Sonifier SFX 150, Emerson). Samples underwent two cycles of sonication in pulses (1s ON, 45ms OFF, for 30s, with 15% intensity), with 1-minute incubation on ice between them. Protein extracts were centrifuged for 10 minutes at 12000g at 4ºC to pellet tissue and cell debris. Supernatant was collected and the total amount of protein was quantified using a Bradford assay (Bio-Rad, USA). All samples were loaded with the same amount of total protein (40μg) and separated on 15% sodium dodecyl sulfate-polyacrylamide gel electrophoresis (SDS-PAGE) and transferred onto PVDF membranes (GE Healthcare, Buckinghamshire, UK) using a Transfer-Blot system (Bio-rad, USA). After blocking with a 5% BSA solution in TBS-T (20mM Tris base, 137mM NaCl and 0.1% Tween-20) for 1 hour, membranes were incubated and cut depending on their molecular weight (KDa) and incubated with primary antibodies overnight at 4°C in a shaker (Cofilin Total 1:500 Invitrogen #MA517275; Phosphorylated-cofilin 1:500 Invitrogen #44-1072G and β-actin 1:3000, Invitrogen #P5-852912). After this period, membranes were washed and then incubated with the corresponding secondary antibodies for 1h at room temperature (Goat Anti-rabbit Secondary Antibody, HRP conjugate 1:1000, ThermoFisher #31460). All antibodies were diluted in a 5% BSA solution in TBS-T. Immunoreactivity was visualized using SuperSignal West Pico PLUS Chemiluminescent Substrate (ThermoFisher, US), using appropriate exposure time in a Chemidoc System (Bio-Rad, USA). The signal intensity was quantified by ImageJ software.

### Data Analysis

Electrophysiological data was collected using a Dagan IX2-700 amplifier (Dagan Corporation, Minnesota, USA) and band-pass filtered (low pass 1 kHz, high pass filter 1 Hz, LHBF 48X, NPI Electronic GmbH, Germany). Data was sampled at 5 kHz using a Lab-PCI-6014 data acquisition board (National Instruments, Austin, TX) and stored on a PC. Offline data analysis was performed using a customized LabView-program (National Instruments). As a measure of synaptic strength, the initial slope of the evoked fEPSPs was calculated and expressed as percent changes from the baseline mean. Error bars denote SEM values. Experiments were rejected if the control pathway decayed more than 20% below baseline. After assessing normality, to test for group differences between LTP values across conditions, an ANOVA was performed with a Tukey post-hoc test (Prism software). For the LTP analysis, fEPSP slope values were averaged in 10 minutes data bins at two-time windows - initial 10 minutes after LTP induction (T1) and at the end of the recorded time (T2=190-200 min or T3=230-240min). The percentage of LTP decay was calculated by [(T1-T2)/T1*100] or [(T1-T3)/T1*100]. For western-blot analysis, cofilin and cofilin-phosphorylated for each sample were normalized to the correspondent β-actin (loading control) and the differences between conditions were assessed by comparing the ratio between cofilin-phosphorylated/cofilin. An ANOVA analysis with a Tukey post-hoc test was performed to assess multiple comparisons.

## Results

### Cdc42 inhibition blocks the maintenance of LTP in a time-dependent manner

Previous studies from our laboratory have shown that interfering with actin cytoskeleton dynamics blocks the induction of persistent forms of synaptic plasticity (Fonseca, 2012; Szabó et al., 2016). Given the known role of Cdc42 in modulating actin dynamics, we sought to assess whether the activation of Cdc42 is necessary for the induction of maintained forms of LTP. To do this, we bath-applied ML141, a selective Cdc42 inhibitor, either at the time of LTP induction or at different time points after LTP induction. We found that inhibiting Cdc42 at the time of LTP induction blocked the maintenance of LTP while blocking Cdc42 at later time points (90-120 minutes) had no impact on LTP maintenance (Figure 1A-E). Inhibition of Cdc42, at an intermediate time point (40 minutes after induction), also had a small destabilizing effect on LTP maintenance although the effect is smaller than the one observed if ML141 is applied at the time of induction (Figure 1B). We also tested the impact of Cdc42 inhibition in transient forms of LTP and found that Cdc42 inhibition has no impact on the expression of transient forms of LTP. In this case, the weak LTP is not altered by Cdc42 inhibition (Figure 1D). As described above, Cdc42 modulates actin dynamics by increasing the phosphorylation of cofilin, inhibiting its activity. To assess if Cdc42 inhibition had an impact on cofilin phosphorylation, we performed a western blot analysis and compared the levels of phosphorylated cofilin in hippocampal CA1 isolated regions. We observed that LTP induction, either by weak or strong synaptic stimulation, significantly increases cofilin phosphorylation and that ML141 application blocks this increase (Figure 1F). This increase in cofilin phosphorylation upon LTP induction is consistent with previous studies Interestingly, weak stimulation led to a similar increase in cofilin phosphorylation as seen for strong stimulation and ML141 application did not affect cofilin phosphorylation in non-stimulated slices. Taken together, our results show that Cdc42 activation is necessary for the maintenance of long-lasting forms of synaptic plasticity in a time-restricted manner. Also, cofilin phosphorylation induced by weak stimulation is similar to cofilin phosphorylation induced by strong stimulation although the expression of LTP is very different. This suggests that Cdc42 activation and downstream modulation of cofilin and actin is a component of the tag and does not seem to be involved in determining the degree of LTP expression.

**Figure 1.**
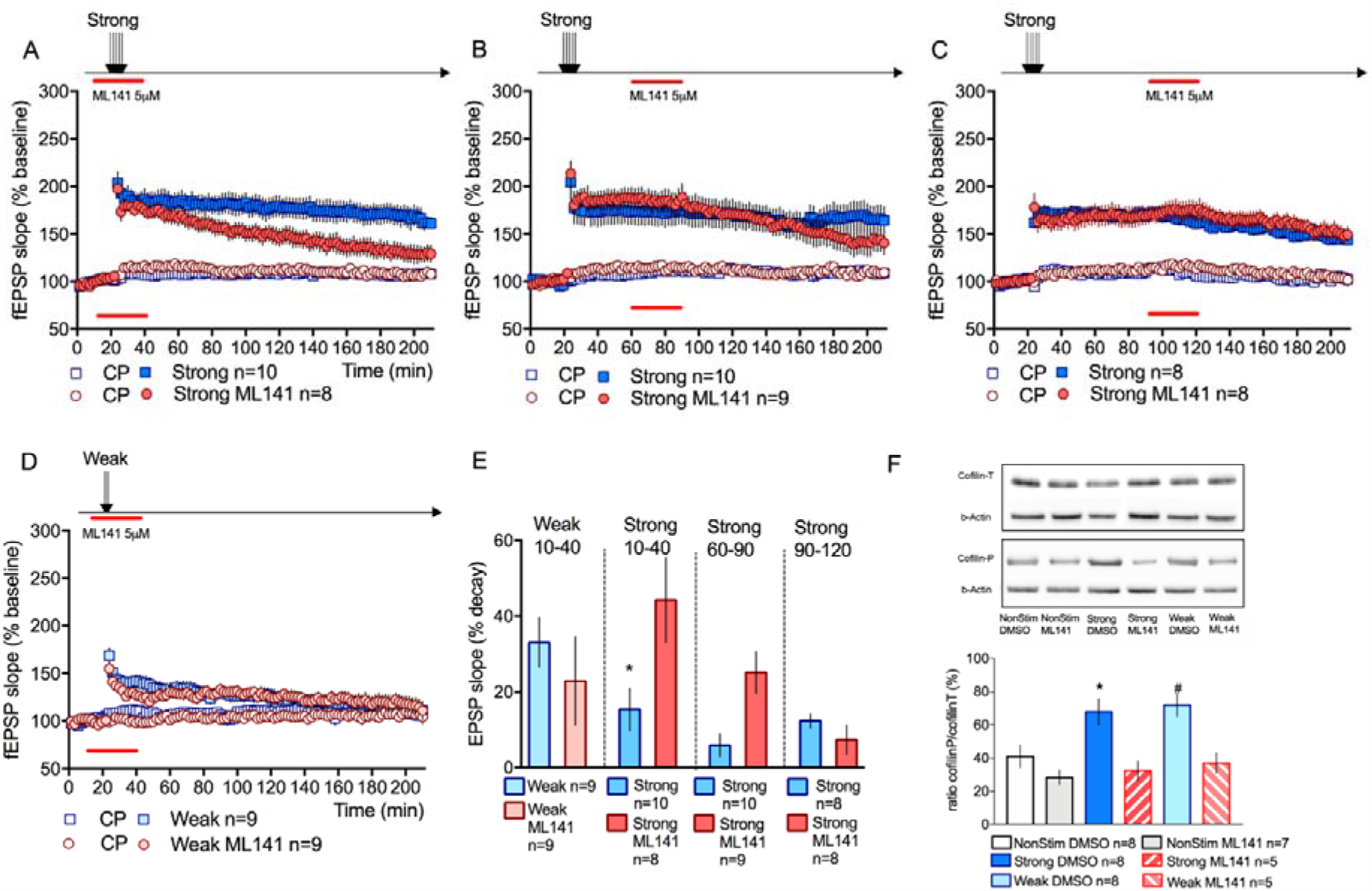
Cdc42 inhibition impairs LTP maintenance. **A**. Inhibiting Cdc42 by bath application of ML141 5μM for 30 minutes during LTP induction blocks the expression of a maintained form of LTP. LTP % decay in ML141 treated slices is significantly higher than in control slices; ML141 44.3±11.1% n=10; DMSO 15.4±5.6%; n =8. **B**. If ML141 is applied at a later time point, from 60 to 90 minutes, LTP maintenance is mildly affected ML141 25.1±5.5% n=9 DMSO 5.9±2.9% n=10 (p=0.054). **C**. Application of ML141 more than one hour after induction (90-120 minutes) has no impact in LTP maintenance ML141 7.4±3.9% n=8 DMSO 12.3±1.9% n=8. **D**. Cdc42 inhibition at the time of a weak LTP induction has no impact in this transient form of LTP ML141 22.9±11.7% n=9 DMSO 33.1±6.5% n=9. **E**. Summary of statistical analysis of experimental conditions. LTP decay is significantly higher when Cdc42 is inhibited at the time of strong LTP induction [F(5,47)=5.89 p=0.0003; *p=0.002] **F**. Western-blot analysis of phosphorylated cofilin in different tested conditions (Total Cofilin (18KDa); Phosphorylated Cofilin (19KDa); b-actin (42KDa). Weak or Strong synaptic stimulation increases the phosphorylation of cofilin, at 30 minutes after induction, which is blocked if Cdc42 is inhibited [F(5,35)=8.08 p<0.001; *p<0.01 #p<0.05; p values correspond to multiple comparisons to all other conditions]

### Cdc42 activation is necessary for the induction of heterosynaptic cooperation

We have previously shown that a dynamic actin cytoskeleton is necessary for heterosynaptic cooperation (Fonseca, 2012). Thus, we expect that inhibiting Cdc42 also blocks synaptic cooperation. Synaptic cooperation results in the stabilization of transient forms of plasticity induced by weak synaptic stimulation (Figure 2B, open symbols), when, within a short time-interval, a second independent pathway is strongly stimulated (Figure 2B, blue symbols). As expected, we found that ML141 application for 30 minutes, between weak and strong synaptic stimulation, blocked the induction of synaptic cooperation preventing the maintenance of the transient LTP induced by weak stimulation of input S1 (Figure 2C). Interestingly, suspending the test pulse stimulation (Figure 2D) or co-application of Cytochalasin 0,5μM (Figure 2E), an inhibitor of actin polymerization, restored synaptic cooperation under Cdc42 inhibition. Group analysis reveals that Cdc42 inhibition significantly increases the decay of weak LTP, blocking the heterosynaptic synaptic cooperative maintenance (Figure 2E). These observations support our hypothesis that Cdc42 activation is involved in setting the synaptic tag that allows synapses to interact by cooperation and that ongoing synaptic activation is an important determinant of actin modulation determining the time-window in which synapses can heterosynaptically interact.

**Figure 2.**
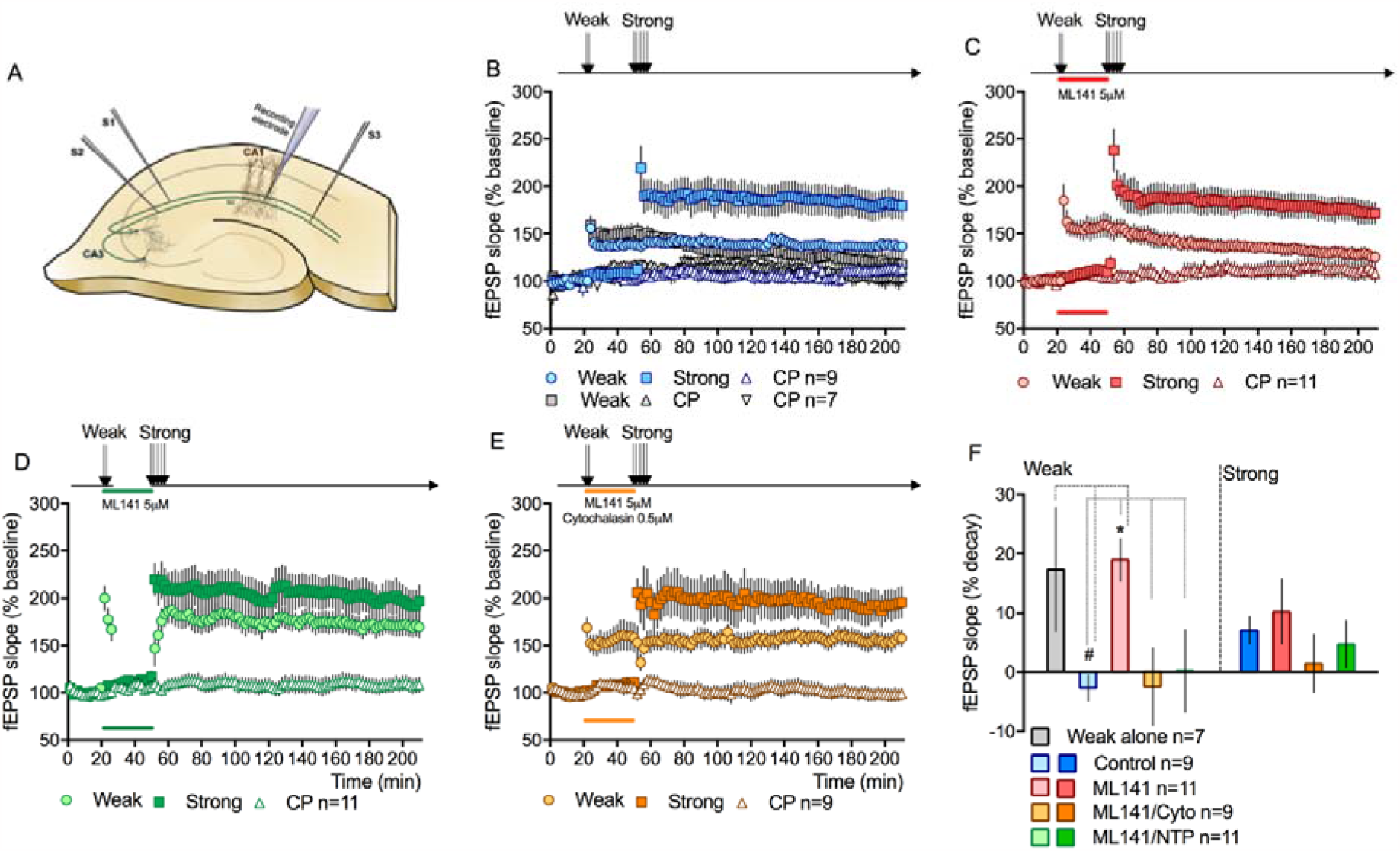
Synaptic cooperation is prevented by Cdc42 inhibition **A**. Schematic representation depicting the position of stimulating and recording electrodes. **B**. Synaptic cooperation: a strong stimulation of S2 stabilizes LTP in the weakly stimulated S1. Percentage decay of LTP in S1 input was significantly lower than in case where weak stimulation is not followed by strong S2 stimulation; Weak alone 17.4±10.5% n=7 Weak cooperation -2.7±2.2% n=9 Strong cooperation 7.04±2.3% n=9. **C**. If ML141 is applied between the weak and strong stimulation (30 minutes of bath application), LTP induced by weak stimulation is no longer maintained. Weak ML141 18.9±3.6% n=11 Strong ML141 10.2±5.4% n=11 **D**. Suspending the test-pulse stimulation of the weakly stimulated S1 input restores its ability to cooperate with Strong S2 even if Cdc42 is inhibited. Weak ML141/NTP 0.18±6.9% n=11 Strong ML141/NTP 4.7±3.9% n=11. **E**. Co-application of cytochalasin with ML141 restores synaptic cooperation. Both drugs were bath applied for 30 minutes between weak and strong stimulation. Weak ML141/Cyto -2.4±6.5% n=9 Strong ML141/Cyto 1.5±4.9% n=9 **F**. Summary of statistical analysis of experimental conditions. LTP decay in Weak S1 is significantly lower in the cooperation setting compared to ML141 treated slices and to the Weak alone experiment [F(4,42)=3.1 p=0.023; #p<0.05]. Weak LTP decay in ML141 treated slices is significantly higher than in control, ML141/NTP and ML141/Cyto conditions *p<0.025.

### Competition is an active heterosynaptic destabilization process

As stated before, synaptic competition is a heterosynaptic form of plasticity that results in the destabilization of LTP maintenance (Fonseca, 2015). The mechanisms behind synaptic competition are even more elusive than for cooperation. We sought to explore whether Cdc42 activation is necessary for synaptic competition. Experimentally, synaptic competition is observed when a third input is activated in close temporal proximity to previously activated inputs, supposedly creating an additional tag that competes with a common pool of available PRPs. Figure 3A illustrates this phenomenon: the activation of a third input S3 leads to the destabilization of the LTP expressed in the Weak S1 and Strong S2. Interestingly, inhibition of Cdc42, by ML141 application during the activation of the Weak S3 input, blocks competition (Figure 3B). All activated inputs expressed maintained forms of LTP, with no significant decay over time. The observation that in this scenario Cdc42 inhibition has no impact on LTP maintenance, contrary to what we observed in our initial experiments (Figure 1A/B), suggests that increasing the number of activated inputs lowers the threshold for LTP induction and expression, rendering it less sensitive to Cdc42 inhibition. Moreover, the observation that inhibiting Cdc42 blocks synaptic competition suggests that competition is a negative heterosynaptic mechanism that actively depresses LTP expression rather than an uneven distribution of resources among different groups of synapses. To address the role of actin dynamics in synaptic competition we use jasplakinolide to inhibit actin depolymerization. Previous results from our laboratory showed that jasplakinolide application blocked synaptic cooperation (Fonseca, 2012). Similarly, we found that stabilizing actin filaments, by jasplakinolide application during the activation of the weak S3 input, blocked competition (Figure 3C) and also blocked LTP expression in the S3 input. Interestingly, we observed that promoting actin depolymerization, by co-applying cytochalasin and ML141, restores the negative interaction between activated synapses, leading to the destabilization of all activated inputs by synaptic competition (Figure 3D). As for cooperation, promoting actin depolymerization counteracts the effect of blocking Cdc42 activation.

**Figure 3.**
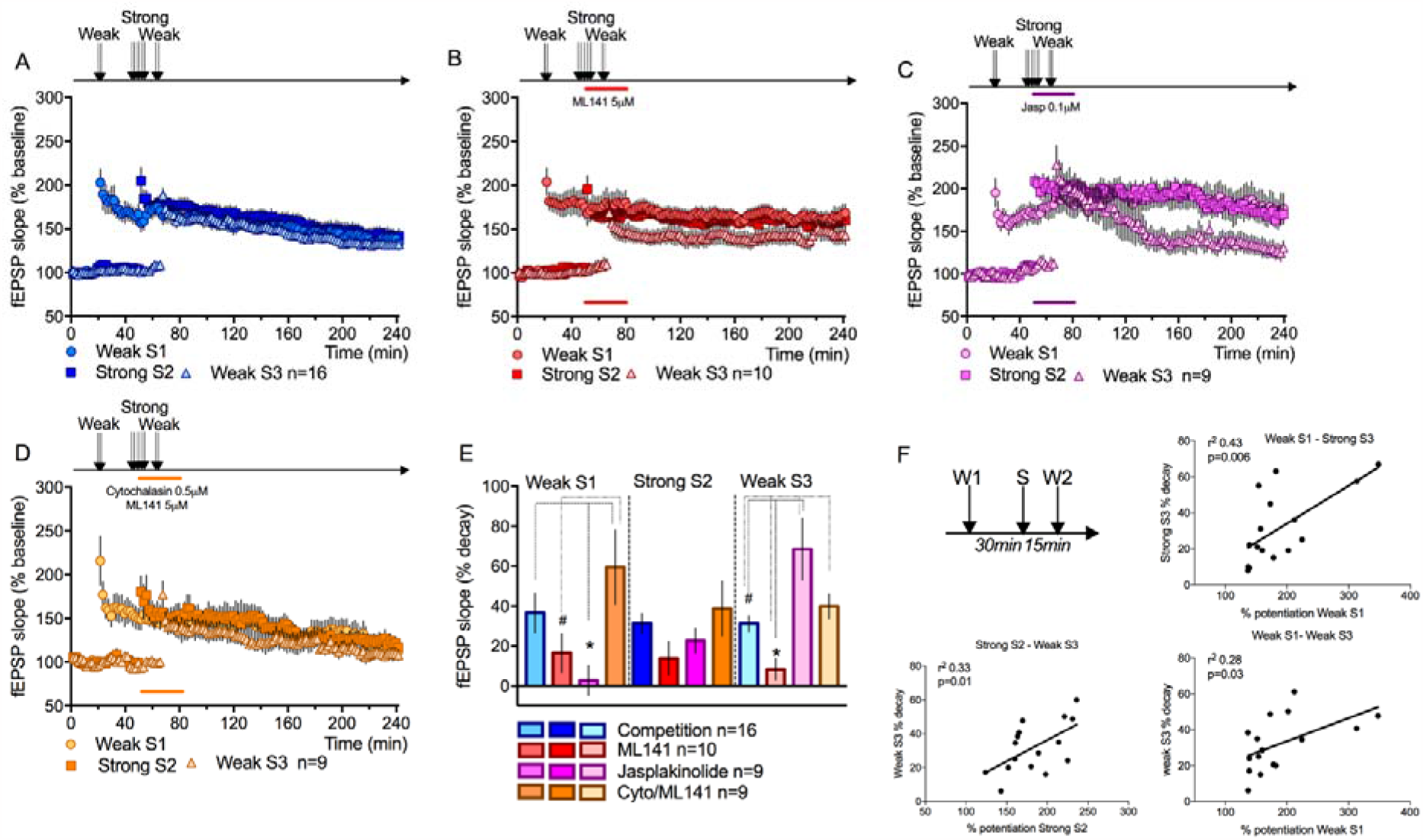
Cdc42 inhibition in a competitive setting promotes cooperative maintenance. **A**. The stimulation of a third input (S3) induces a competitive setting leading to LTP decay in all activated inputs. LTP percentage decay Weak S1 36.5±9.8% Strong S2 31.2±5% Weak S3 31.1±4% n =16. **B**. If ML141 is applied, for 30 minutes after strong S2 stimulation, competition is blocked and converted in cooperation. LTP percentage decay Weak S1 16.5±9.7% Strong S2 13.8±8.4% Weak S3 8.3±5.5% n=10 **C**. Stabilizing actin filaments by jasplakinolide application blocks competition but prevents the stabilization of LTP in the Weak S3 input. Weak S1 2.7±7.4% Strong S2 22.5±6.5% Weak S3 68.5±15.5% n=9 **D**. Co-application of cytochalasin with ML141 restores competition resulting in the destabilization of LTP in all activated inputs. Weak S1 59.4±18.5% Strong S2 38.6±13.8% Weak S3 39.9±6.1% n=9 **E**. Summary of statistical analysis of experimental conditions. LTP decay is not significantly different for the strong S2 input in all conditions tested [F(3,40)=1.59 p=0.20]. For Weak S1, LTP decayed significantly less in Jasplakinolide-treated slices as compared to control and ML141/cyto conditions [F(3,40)=3.75 p=0.01; #p<0.05] and in ML141 as compared to ML141/cyto treated slices (*p=0.02). For Weak S2, LTP decayed significantly less in ML141 treated slices as compared to all other conditions [F(3,40)=8.84 p<0.001; *p<0.04] and significantly more in Jasplakinolide treated slices as compared to control (#p=0.001). **F**. Competition has a time rule. The percentage decay of LTP in strong S2 and Weak S3 is statistically correlated with the percentage potentiation of W1.

One interesting aspect of heterosynaptic competition is the observation that its induction depends on the order and the temporal relationship between activated synaptic inputs (Madeira et al., 2020). To illustrate this temporal rule, we plotted the percentage decay for each activated input, Weak1, Weak2 and Strong in relation to the percentage potentiation of each input and found statistically significant correlations between sequentially activated inputs (Figure 3F). That is, we observe that the percentage decay of the Weak2 is correlated to the percentage potentiation of the Weak 1 or the Strong input and that the percentage decay of the Strong is correlated to the percentage potentiation of the Weak1 (Figure 3F). This temporal correlation decreases by increasing the time interval between stimulations (Weak1-Weak2) and there is no correlation between reverse interactions (data not shown). This observation suggests that the degree of competition is determined by previous activity in a time-dependent manner. Taken together, our results support the hypothesis that an active actin dynamic cytoskeleton is necessary for both heterosynaptic interactions, synaptic cooperation as a positive re-enforcement of LTP maintenance and synaptic competition, a negative interference of LTP maintenance (Figure 3E).

### Heterosynaptic cooperation and competition also occur between weak synapses

As stated above, the STC hypothesis proposes that synaptic cooperation and competition are based on the distribution of PRPs among activated synapses. In this scenario, synapses engage in cooperation or competition depending on the availability of PRPs in relation to the number of activated synapses. Also, it is assumed that weak synaptic stimulation does not induce the synthesis of PRPs given that changes in synaptic efficacy induced by this stimulation are transient and insensitive to protein synthesis inhibitors. This idea that weak stimulation does not induce the synthesis of proteins is even more important in the case of competition given that in this situation the observed destabilization in all activated inputs is attributed to an uneven distribution of PRPs produced by the strong stimulation of one input and subsequent distribution of PRPs among all three inputs (two weak and one strong). If conversely, cooperation and competition are heterosynaptic forms of potentiation and depression then one should be able to induce cooperation and competition between weakly stimulated inputs. To test this hypothesis, we designed a cooperation experiment where the second input also received a weak stimulation. We observed that the two inputs engage in cooperation and LTP is maintained in both stimulated inputs (Figure 4A). It is noteworthy to mention, that if the second input S2 is not stimulated, the weak stimulation in S1 results in a transient LTP that returns to baseline (Figure 4B). Interestingly, if a third input is further stimulated, with a weak stimulation, synapses engage in competition, resulting in the destabilization of LTP in all activated inputs (Figure 4C). These results are similar to what we observed previously when S2 was strongly stimulated (Figure 3A). Taken together, our results show that synaptic cooperation and competition can be induced between inputs that are weakly stimulated suggesting that these heterosynaptic interactions are more than just the capture of PRPs at activated synapses.

**Figure 4.**
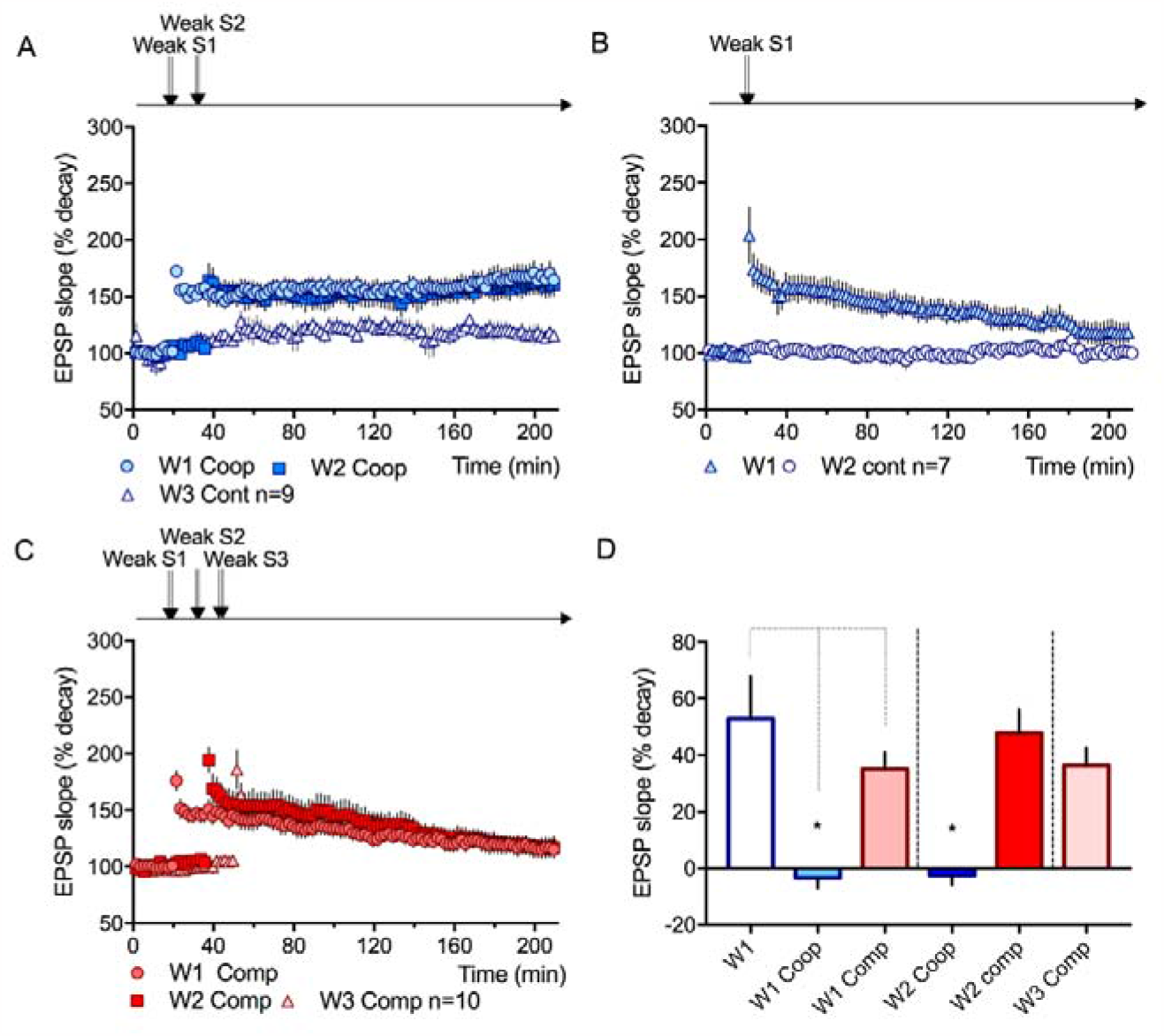
Cooperation and competition are heterosynaptic facilitation and depression mechanisms, respectively. **A**. Cooperation was induced by stimulating two inputs with a weak stimulation. Percentage LTP decay Weak S1 -3.4±3.5% Weak S2 -2.43±3.3% n =9. **B**. Weak stimulation alone is not sufficient to induce a maintained form of LTP. Percentage decay of LTP Weak S1 52.8±15.03% n=7 **C**. The stimulation of a third weak input induces competition resulting in the destabilization of all stimulated inputs. Weak S1 35.1±5.9% Weak S2 47.8±8.5% Weak S3 36.3±6.3% n=10. **D**. Summary of statistical analysis of experimental conditions. For the Weak S1 input, LTP decay is significantly lower in the cooperation setting compared to the weak alone or the competition setting [F(2,23)=11.6 p<0.001; *p<0.01]. Similarly, for the Weak S2 input, LTP decayed significantly less in the cooperation setting as compared to the competitive setting (Unpaired T-test, *p<0.01).

## Discussion

Synaptic cooperation and competition are two forms of heterosynaptic plasticity that allow neurons to integrate activity across time and within their extensive dendritic tree (Fonseca, 2015). Heterosynaptic forms of plasticity have been suggested to be an important mechanisms to scale and normalize synaptic weights (Chistiakova et al., 2014), preventing saturation of synaptic responses and runway by concurrent neuronal activity. Thus, heterosynaptic plasticity effectively promotes the stabilization of homosynaptic plasticity, while balancing the overall excitability of the activated neuron (Field et al., 2020). The key difference between synaptic cooperation and competition, described here, from other forms of heterosynaptic plasticity is their time course. Classic heterosynaptic LTP (or LTD) is observed at the time of induction of homosynaptic plasticity and is based on the spreading of calcium within dendrites (Chistiakova et al., 2014). Conversely, the forms of heterosynaptic plasticity described here can be observed within relatively high time intervals (30 to 60 minutes) and are thought to be based on two independent mechanisms, the setting of an input-specific synaptic tag and the synthesis of plasticity-related proteins (PRPs) that are captured by the tags (Frey and Morris, 1997; Redondo et al., 2010). Given that the tag is responsible for the capture of PRPs, the time window in which synaptic cooperation and competition can be observed depends on the time in which the tag is active after the induction of synaptic plasticity. Thus, comprehending the nature of the tag and its dynamics is crucial to understanding the time frame of such heterosynaptic interactions and ultimately how these mechanisms contribute to memory acquisition. We have previously proposed that the tag, rather than a specific molecule, is a state of the synapse that in an activity and time-dependent manner renders synapses permissive to change (Madeira et al., 2020; Pinho et al., 2020; Szabó et al., 2016).

Here, we assessed the role of a Cdc42, a well-known Rho GTPase and regulator of actin dynamics, as a component of the synaptic tag, modulating the maintenance of homo and heterosynaptic forms of synaptic plasticity. We used ML141, a non-competitive inhibitor, highly selective for Cdc42 (Surviladze et al., 2010) that acts by blocking the binding of Cdc42 GTPase to guanine nucleotides (Hong et al., 2013). We show that inhibition of Cdc42 at the time of LTP induction blocked the maintenance of LTP and that this blockade occurred in a time-limited manner, given that Cdc42 inhibition had no impact on LTP maintenance if ML141 was applied 1 hour after LTP induction. These results suggest that Cdc42 is important during the initial alterations in synapses that allow, later on, the maintenance of LTP. It is likely, that Cdc42, RhoA and Rac1 are all necessary for functional and structural maintenance of LTP and that their activation complements each other to transduce extracellular signals that, through activation of NMDA receptors, BDNF and CaMKII signaling result in synaptic remodeling and strengthening of synapses (Duman and Tolias, 2016; Hedrick et al., 2016; Hedrick and Yasuda, 2017). However, inhibiting Cdc42 is sufficient to block LTP maintenance. Consistent with the role of Cdc42 in modulating cofilin phosphorylation, we found that ML141 application inhibited the increase in cofilin phosphorylation observed upon the induction of LTP. It is noteworthy to mention that we observed that both weak and strong synaptic stimulation leads to a similar increase in cofilin phosphorylation and that ML141 application resulted in a similar decrease in cofilin phosphorylation in both experimental conditions. This contrasts with the observation that ML141 application did not affect the expression or maintenance of LTP induced by weak synaptic stimulation. Combined, these results support our hypothesis that Cdc42 activation is part of a cellular mechanism that tags activated synapses but is not directly involved in determining the degree of plasticity expressed.

Previous studies have shown that structural and molecular remodeling of individual dendritic spines upon LTP was divided into two temporal phases (Bosch et al., 2014). An initial phase consisted of an actin cytoskeleton remodeling phase (<7 min), accompanied by a rapid increase of actin, Arp2/3 and cofilin concentration in the spine. The authors postulate that cofilin and Arp2/3 (both downstream of Cdc42) might act synergistically to remodel actin filaments, creating new docking sites for plasticity-related proteins. At the same time, the concentration of proteins that could stabilize and compete with cofilin and Arp2/3 for F-actin binding sites is transiently reduced (e.g. CaMKIIβ, drebrin) (Borovac et al., 2018; Hedrick and Yasuda, 2017; Kim et al., 2016; Murakoshi et al., 2011). Thus, decreasing actin-stabilizing proteins and increasing cofilin concentration in synapses contribute to active actin filament remodeling. The second, stabilization phase (>7 min) was associated with the formation of a stable complex of cofilin with F-actin. Cofilin was persistently retained at potentiated spines for more than 30 minutes, which fits very well with the time window of synaptic cooperation and competition. Interestingly, cofilin activity has been described as being dependent on concentration (Andrianantoandro and Pollard, 2006). At a low stoichiometric ratio of cofilin, single cofilins bind and sever actin filaments while in higher concentrations, cofilin binds cooperatively to F-actin, promoting actin filament assembly (Andrianantoandro and Pollard, 2006). Thus, cofilin has a bidirectional effect on actin dynamics and is a necessary protein for the consolidation of functional and structural LTP (Bosch et al., 2014; Rust, 2015). As such, cofilin must be inactivated in time, not only to be retained in the spine but also to avoid reverting the polymerizing trend toward depolymerization during the stabilization phase. A decrease in cofilin phosphorylation by Cdc42 inhibition results in a blockade of this stabilization phase, preventing LTP maintenance.

Consistent with this role of Cdc42 in modulating actin dynamics, we found that inhibiting Cdc42 blocked synaptic cooperation. Although ML141 application had no effect on the initial expression of LTP induced by a weak synaptic stimulation at input S1, Cdc42 inhibition blocked the cooperative maintenance of LTP induced by the subsequent strong S2 synaptic stimulation. This result is consistent with our previous observations in which directly interfering with the actin cytoskeleton dynamics also blocked synaptic cooperation (Fonseca, 2012). Interestingly, we found that synaptic capture is highly influenced by ongoing neuronal activity which supports our hypothesis that synaptic activation, after LTP induction, modulates the time window in which the tag is active and thus, the time window in which synapses can interact by cooperation. In other words, synaptic activation closes the permissive state of synapses. By suspending ongoing synaptic activation, we were able to prolong the time window of cooperation reverting the effect of Cdc42 inhibition. We have also observed this modulatory effect of synaptic activity in the time window of synaptic cooperation in the amygdala circuitry (Fonseca, 2013; Madeira et al., 2020), showing that excitatory synapses of pyramidal neurons have similar temporal dynamics. The blockade of Cdc42 inhibition in synaptic cooperation was also reverted by promoting actin depolymerization, suggesting that increasing G-actin availability is able to counteract the inhibition of Cdc42. Although at first glance this result may look contradictory, there is strong evidence that different pools of actin filaments have different dynamics and are differentially affected by pharmacological inhibitors such as cytochalasin B (Duman and Tolias, 2016; Shekhar and Carlier, 2017; Skruber et al., 2018; Sumi et al., 1999; Wioland et al., 2017). In our experimental condition, the use of cytochalasin B results in a net increase in G-actin availability, which has been shown to promote a switch in cofilin activity towards polymerization of actin filaments.

Cdc42 activation is also required for synaptic competition. As described previously, inducing LTP in a third input with a weak stimulation leads to the destabilization of LTP in the two previously activated inputs, that is, heterosynaptic competition. Cdc42 activation inhibition blocked competition consistent with our hypothesis that a permissive state of the synapse, due to actin modulation, determines whether synapses interact or not. As for cooperation, Cdc42 inhibition prevents synapses from progressing to the permissive state. This is also supported by the observation that promoting actin depolymerization restores competition. The observation that, under competition, Cdc42 inhibition allows the maintenance of LTP in all activated inputs shows that competition is a heterosynaptic destabilizing process, in which activated synapses actively promote the depotentiation of other synapses, rather than the distribution of a limited pool of PRPs among tagged synapses. If synaptic competition is a form of heterosynaptic depotentiation or heterosynaptic LTD, why is it necessary to activate three inputs to observe this negative interaction? Heterosynaptic interactions are typically observed in a doughnut shape, with synapses closed to the potentiated ones showing heterosynaptic LTP and more distal ones showing heterosynaptic LTD (Chistiakova et al., 2014; Volgushev et al., 2016). We hypothesise that by recruiting more synapses, as more inputs are stimulated, the overall excitability of the neuronal networks also increases, in turn increasing this doughnut shape effect and promoting an increase of the LTD area. Supporting this, yet speculative hypothesis is the observation that in the competitive setting, the degree of potentiation of the first stimulated input is correlated to the degree of LTP decay in the subsequent activated inputs. In other words, the more potentiated the first input is, the higher the decay in the subsequent activated inputs. If the degree of potentiation alters the shape of the heterosynaptic doughnut, then the higher potentiation of the first stimulation the higher the probability that the next stimulation leads to heterosynaptic depression.

As stated in the introduction, in light of the synaptic tagging and capture model (STC), it has been proposed that synaptic cooperation and competition are based on the interplay between synaptic-specific tags and the capture of PRPs among tagged synapses. This hypothesis is substantiated by previous studies showing that synaptic cooperation and competition are sensitive to protein synthesis inhibitors and it assumes that weak stimulation generates tags but not PRPs. We propose that synaptic cooperation and competition are due to changes in neuronal excitability, that alter the threshold of heterosynaptic plasticity. Depending on the number of inputs that are stimulated, the increase in excitability favours heterosynaptic LTP, or cooperation, and if a third input is stimulated the further increase in excitability favours heterosynaptic LTD. This hypothesis is supported by our data showing that synaptic cooperation and competition can be observed if two or three inputs are weakly stimulated. If one considers the STC model, then cooperation should not be observed when two inputs are weakly stimulated since neither should have reached the threshold for PRPs synthesis. If rather, weak stimulation increases excitability, then the threshold for LTP induction is lower when the second input is weakly stimulated leading to the cooperative effect of LTP maintenance in both inputs. In competition, the stimulation of the third input results in heterosynaptic LTD. It is noteworthy to mention that several PRPs, involved in LTP maintenance and synthesized upon LTP induction are regulators of neuronal excitability, such as Homer, BDNF, PKA and PKMζ (Desai et al., 1999; Zhang et al., 2012; Zhong et al., 2009). Thus, protein synthesis inhibition results in a decrease in neuronal excitability which would explain why synaptic cooperation is blocked by protein synthesis inhibition.

Overall, our results support the hypothesis that, by regulating the actin cytoskeleton, Cdc42 interferes with the maintenance of homo and heterosynaptic forms of synaptic plasticity. Actin cytoskeleton remodeling as a consequence of neuronal activity renders synapses in a permissive state that allows the synaptic composition to change, in a time-dependent manner. While in this permissive state, synapses can interact positively, cooperating to maintain plasticity, or negatively, interfering with the maintenance of synaptic plasticity.

## Acknowledgements

We would like to thank the histology facility and the animal facility at NOVA Medical School for the implementation of the electrophysiology and molecular biology experiments. This work was funded by a Portuguese Research Council grant (Fundação para a Ciência e Tecnologia - PTDC/NEU-SCC/3023/2014) and a CEEC grant to Rosalina Fonseca (2020.02221.CEECIND).

## Notes

### Competing Interest Statement

The authors have declared no competing interest.

